## Supplemental Information for "gUMI-BEAR, a modular, unsupervised population barcoding method to track variants and evolution at high resolution"

Supplementary information

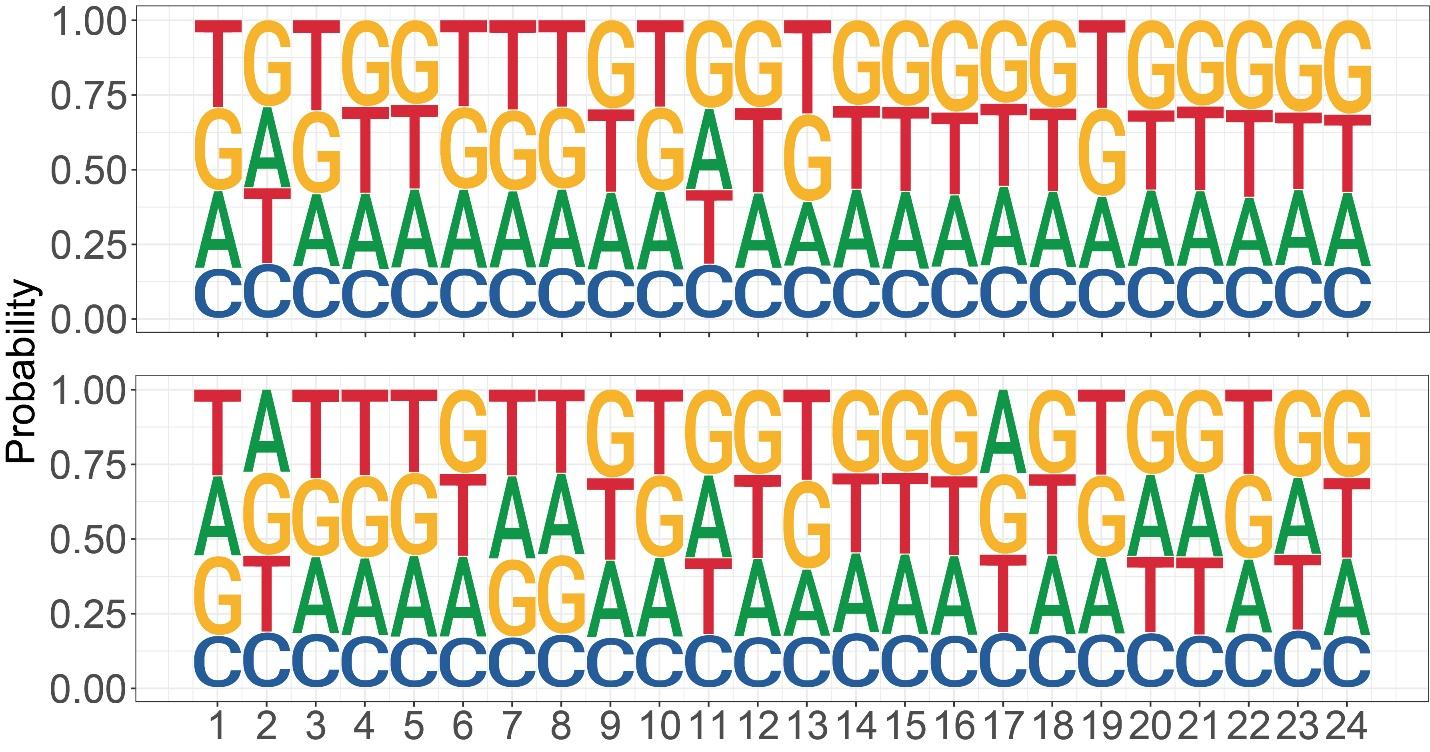

**Supplementary Figure 1. Nucleotide composition by position for the gUMI from donor DNA and lineages**

Donor DNA prior to transformation (upper panel) and DNA from lineages participating in the experiment (lower panel) were extracted and deep-sequenced to reveal the nucleotide composition for each position in the gUMI (x-axis). The overall height of each base is proportional to its probability (y-axis). Colours represent different nucleotides.

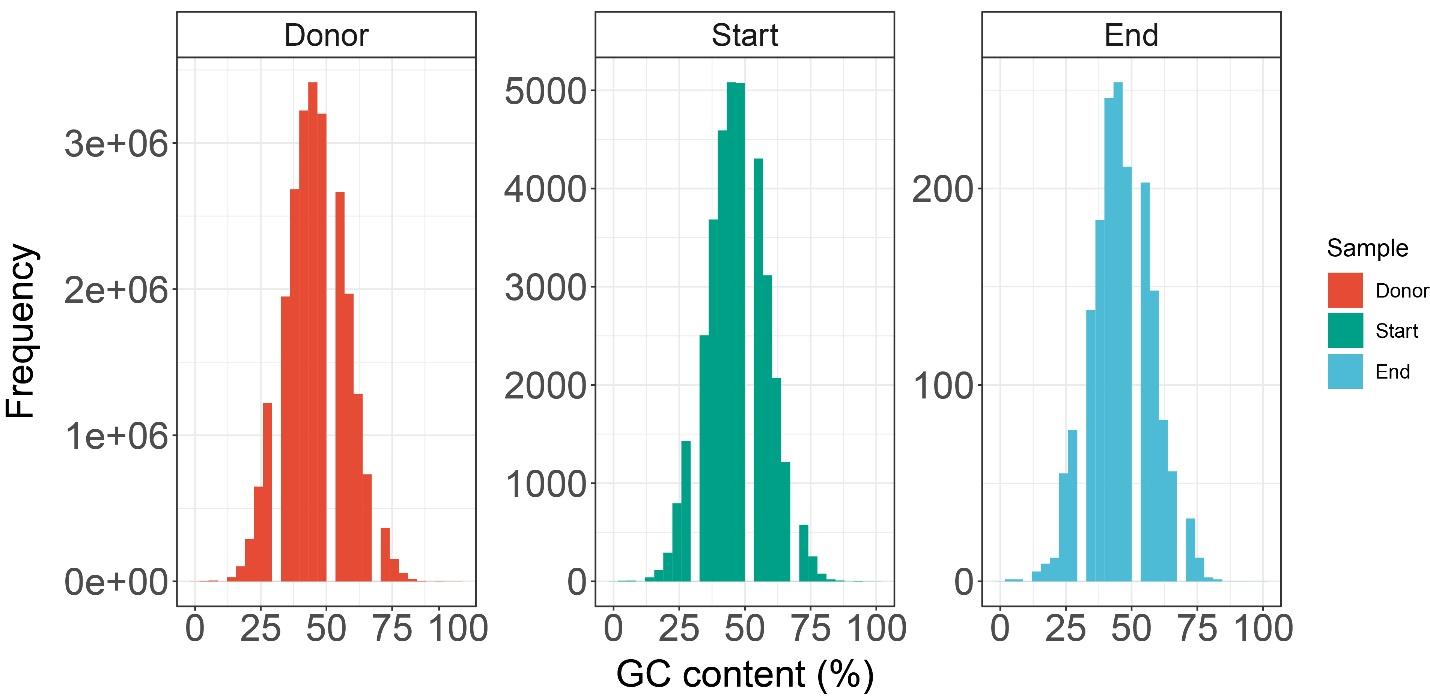

**Supplementary Figure 2. Guanine-Cytosine content distributions for the gUMI from three stages in the experiment with no initial fitness variations**

GC content was measured based on sequences derived from all lineages at three steps during library construction. (Left, red) Donor DNA prior to transformation. (Middle, green) Lineages at the beginning of the experiment. (Right, blue) Lineages at the end of the experiment. All data are presented in histograms to reveal the GC content distribution in the populations.

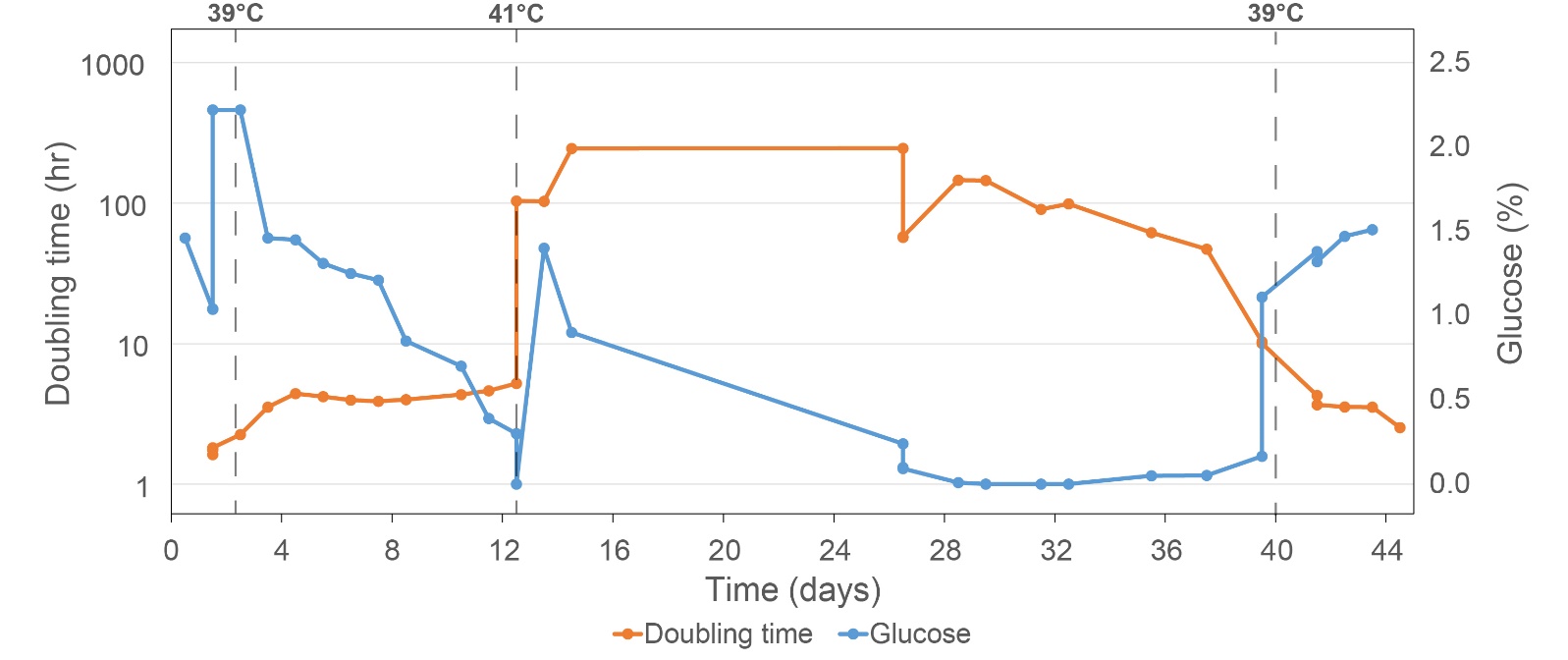

**Supplementary Figure 3. Mean doubling time and glucose concentration for the experiment in population exhibiting no initial fitness variations**

Doubling time for the whole culture was calculated based on dilution rate of the turbidostats and is presented on a log scale (left y-axis, orange line) and ploted as function of time. Glucose concentrations were measured for each sample taken (right y-axis, blue line). The experiment started at an optimal growth temperature of 30°C that was altered to induce stress as noted by the doted lines.

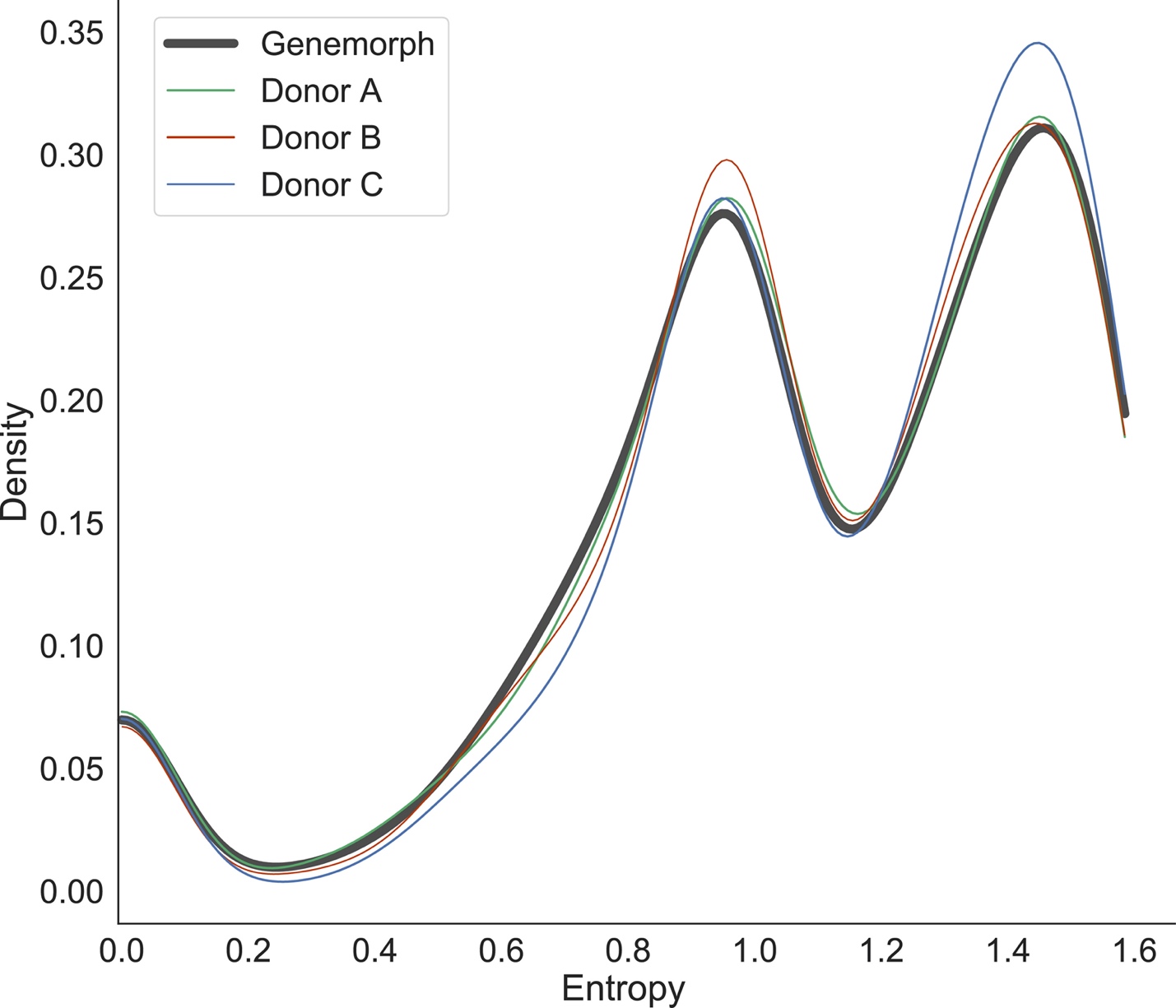

**Supplementary Figure 4. Entropy kernel density estimations for the *HSP82* ORF before and after incorporation into the gUMI-box**

The kernel density estimations of the Shannon entropy for nucleotide substitutions in the *HSP82* ORF in the genenmorph amplicon after *in vitro* mutagenesis (black) and three replicates of the assembled donor DNA (green, red and blue) are shown. Three peaks can be observed - first at entropy 0, in which all substations were to the same nucleotide, second at entropy 1, in which substitutions were equal between one of two possible substitutions, and finally, the third peak at entropy 1.5 in which an equal probability for all mutations is observed. The donor DNA distribution mimics the one produced by the genemorph, indicating that no bias was induced during donor construction.

| Mini-library Name | No. transformations | T_sel_ (h) | No. cells sorted (×10^3^) | T_exp_ (h) | No. unique lineages |
| --- | --- | --- | --- | --- | --- |
| B2 | 10 | 20 | 55 | 31 | 5349 |
| B3 | 10 | 20 | 55 | 31 | 4865 |
| B10 | 10 | 20 | 10 | 31 | 1636 |
| C55 | 1 | 20 | 55 | 31 | 2768 |
| C10 | 1 | 20 | 10 | 31 | 1538 |
| D55 | 10 | 21 | 55 | 34 | 5189 |
| D110 | 10 | 21 | 110 | 34 | 9028 |

**Supplementary Table 1. Mini-library parameters**

Seven different mini-libraries were prepared while varying the following parameters: No. transformations - the factor applied to the volumes given in the protocol published by Ryan et al.^1^, namely, 1 µg pCAS, 5 µg donor DNA, and 90 µL competent cells. Hence, No. of transformations = 10 indicates the use of 10 µg pCAS, 50 µg donor DNA, and 900 µL competent cells; T_sel_, selection time, being the number of hours the library was grown for after transformation and before sorting; No. cells sorted, total number (in thousands) of viable cells sorted following staining with propidium iodide and selection (see Methods); T_exp_, expansion time, being the time (hours) the library was allowed to expand before division into 100 aliquots; No. unique lineages, number of viable lineages in each mini-library as determined by deep-sequencing.

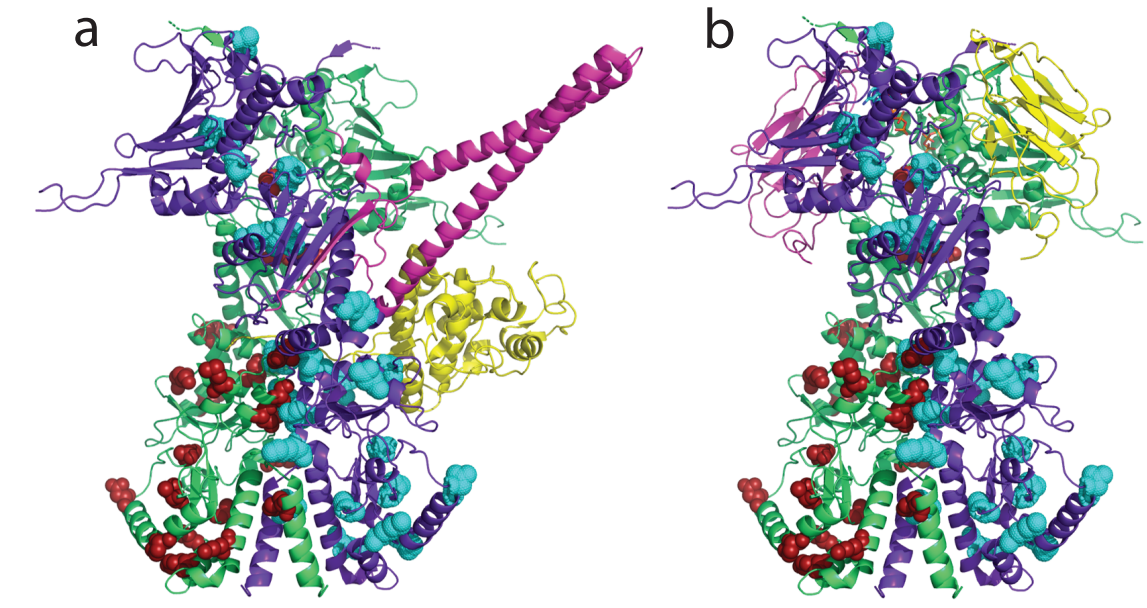

**Supplementary Figure 5. Protein-protein interaction models**

**a**, HSP82 dimer shown in green and purple, Cdc37 cochaperone (pink), and the Cdk4 kinase (yellow) (PDB 5FWK). **b,** HSP82 dimer shown in green and purple and Sba1 Chaperone (in yellow and pink) (PDB 2CG9). All possible mutations are shown in the HSP82 model as red and cyans balls for the two HSP82 homodimer.

**Supplementary results**

**Proteomics analysis of hsp82 variants**

Proteins often carry out their functions through interactions with other proteins to form multi-protein complexes. Such protein-protein interactions play a central role in various aspects of the structural and functional organization of the cell, and their elucidation is crucial for a better understanding of biological processes.

To elucidate the role of the mutations found in the final population in our experiments in their ability to overcome other variations, we analysed their involvment in the dimer interface and in the binding interface with other proteins. The dimer interface of the Hsp82 structure contains 228 contacts up to 3.5 Å involving 42 amino acid residues, of which only five were mutated in five different variants in the final population in our experiments;

N377I (HSP-40), I505V (HSP-11), R591K (GSP-7), A595V (HSP-4) and L598M (HSP-29). It seems reasonable to assume that a single point mutation in a residue involving the dimer interface will interfere with dimer formation. To understand the possible role of the mutations in binding different proteins we modeled these Hsp82 mutations into known structures of Hsp82 with other proteins. Specifically, we modeled the Hsp82 dimer (green and purple in Supplementary Figure 5) into the structure of Cdc37 (pink in Supplementary Figure 5a) cochaperone, Cdk4 kinase (yellow in Supplementary Figure 5a), and Sba1 cochaperone (yellow and pink in Supplementary Figure 5b). None of the 39 mutations we found are involved in heterodimer formation.

**Supplementary methods**

**Restriction Free (RF) cloning and guide-RNA design**

For both applications of our method, we used Chop-Chop^2^ (CHOPCHOP (uib.no)) to find the best matching sequence in our locus of interest for Crispr/Cas9 double-strand breakage.

For the experiment with the population exhibiting no initial fitness differences, we made a single double-strand breakage using the following gRNA sequence:

5’-AGAGCGTCAATCAAGAAAG-3’

and the resulting primers for RF cloning to the pCAS vector:

5’-CGGGTGGCGAATGGGACTTTTAGAGCGTCAATCAAGAAAGGTTTTAGAGCTAGAAATAGC-3’

5’-GCTATTTCTAGCTCTAAAACCTTTCTTGATTGACGCTCTAAAAGTCCCATTCGCCACCCG-3’

For the use with tracking *HSP82* gene variants, we induced two double-strand breaks to integrate our donor DNA into the locus of interest. Two pCAS vectors were cloned using the following gRNA sequences:

Upstream to the *hsp82* gene: 5’-CAAACAAACACGCAAAGATA-3`

Downstream to the *hsp82* gene: 5’-AGCTGACACCGAAATGGAAG-3'

and the resulting primers for RF cloning to the pCAS vector:

Upstream of the *hsp82* gene:

5’-CGGGTGGCGAATGGGACTTTTCAAACAAACACGCAAAGATAAAAGGTTTTAGAGCTAGAAATAGC-3’

5’-GCTATTTCTAGCTCTAAAACTATCTTTGCGTGTTTGTTTGAAAAGTCCCATTCGCCACCCG-3’

Downstream of the *hsp82* gene:

5’-CGGGTGGCGAATGGGACTTTTAGCTGACACCGAAATGGAAGAAAGGTTTTAGAGCTAGAAATAGC-3’

5’-GCTATTTCTAGCTCTAAAACCTTCCATTTCGGTGTCAGCTAAAAGTCCCATTCGCCACCCG-3’

**Donor construction for tracking *HSP82* variants**

The donor DNA construct was assembled in stages, via six PCR reactions that constructed and then assembled two sub-constructs as described in the “Results” section. To build the first sub-construct, the Hsp82 gene was amplified from the genome of the BY4741 yeast strain (forward primer 100 bp upstream of gene - Genemorph_F; reverse primer 100 bp downstream of gene - Genemorph_R ; PCR 1: kappa 50 μL, annealing at 60 °C, 25 cycles, 2 min elongation time, 90 ng genomic DNA template). Random mutations were inserted using the GeneMorph II random mutagenesis kit using the Genemorph_F and Genemorph_R primers (1000 ng of template amplicon and 30 cycles to ensure a low mutation rate of 0–4.5 mutations/kb, PCR 2: annealing at 60 °C, 4 min elongation time). Two overhangs were added to the mutated gene by a single PCR reaction. One overhang (LHA primer) contained a 75 bp sequence upstream and a 5 bp sequence downstream of the integration site. The second overhang (HSP82 _Read1) comprised a 15 bp sequence upstream of the stop codon of the genomic Hsp82 gene and a Read-1 binding site for Illumina NGS platforms (PCR 3: kappa 50 μL, annealing at 60 °C, 25 cycles, 2 min elongation time, 40 ng mutated hsp82 template).

The second sub-construct was made by adding two overhangs. One primer (Gumi2Hyg) contained: a binding site for Read-1; not G nucleotide (5×H); the gUMI barcode comprised of a 24 bp random sequence, a Linker sequence, and a 25 bp sequence complementary to the Hygromycin B resistance cassette as found on the vector pAG32. The second primer (RHA) contained a 70 bp sequence downstream of the stop codon site of the ghsp82 gene and a 26 bp sequence upstream of the stop codon of the Hygromycin B cassette (PCR 4: kappa 50 μL, annealing 60 °C, 25 cycles, 2 min elongation time, 10 ng pAG32 template).

Following PCR cleanup, the full donor DNA construct was assembled by overlapping the two sub-constructs at their Read-1 regions in a PCR reaction that used PrimeSTAR GXL DNA Polymerase (PCR 5: 50 μL annealing at 60 °C, 15 cycles, 4 min elongation time, 10 ng of each construct, no primers). The donor DNA construct was then amplified using a 5 µL aliquot of the product of PCR 5 (primers, LHA & RHA; PCR 6: kappa 50 μL, annealing at 60 °C, 15 cycles, 4 min elongation time). Primers were obtained from IDT (Israel), with the exception of the RHA primer, which that was obtained from Sigma (Israel).

**All primer sequences used for the donor construction process**

**No initial fitness differences experiment**

Gumi2Hyg******

5’-AAATAGGGGAATGAACGCATATTGGTTTCATTATAGAGCGTCAATCAAGATCGTCGGCAGCGTCAGATGTGTATAAGAGACAGHHHHHNNNNNNNNNNNNNNNNNNNNNNNNTTGGAAGTGTGGCTAGACATGGAGGCCCAGAATACCC - 3’

LHA:

5’ – AGCGTTCCTAGCCCTACCGAGAAATGTGCGTTTATAGTTTGGTGTCTCTTCAGTATAGCGACCAGCATTCACAT - 3’

***HSP82* gene variants**

Genemorph_F

5’ - GTGACCTCCTCATTTCTTCCCG - 3’

Genemorph_R

5’- GTGACCTCCTCATTTCTTCCCG - 3’

LHA

5’ - TGTATTAGAGTTCAAGAAATCATACCTGATAGAAAATAGAGTCCTATAAACAAAAGCACAAACAAACACGCAAAGATATG - 3’

HSP82 _Read1

5’ - CTGTCTCTTATACACATCTGACGCTGCCGACGACTAATCTACCTCTTC - 3’

Gumi2Hyg 5’ –TCGTCGGCAGCGTCAGATGTGTATAAGAGACAGHHHHHNNNNNNNNNNNNNNNNNNNNNNNNTTGGAAGTGTGGCTAGACATGGAGGCCCAGAATACCCTCC - 3’

RHA 5’ –TTATTCATTCGAATACCTATACGTTATATTATGTTTTGTTTATAACCTATTCAAGGCCATGATGTTCTACCAGTATAGCGACCAGCATTCACATAC - 3’

**Supplamentary References**

1.	Ryan, O. W., Poddar, S. & Cate, J. H. D. Crispr–cas9 genome engineering in Saccharomyces cerevisiae cells. *Cold Spring Harb. Protoc.* **2016,** 525–533 (2016).

2.	Montague, T. G., Cruz, J. M., Gagnon, J. A., Church, G. M. & Valen, E. CHOPCHOP: A CRISPR/Cas9 and TALEN web tool for genome editing. *Nucleic Acids Res.* **42,** 401–407 (2014).
